# fMRI Network Features Identify Speech and Language Critical Cortex

**DOI:** 10.64898/2026.07.23.740155

**Authors:** Birsu Baç, Robert D. Flint, Zachary Fitzgerald, Jason K. Hsieh, Nathan E. Crone, Melissa-Ann Mackie, Todd B. Parrish, Matthew C. Tate, Richard Betzel, Marc W. Slutzky

**Affiliations:** Department of Neurology, Northwestern University; Department of Neurosurgery, Northwestern University; Department of Neurology, Johns Hopkins University; Department of Psychiatry & Behavioral Sciences, Northwestern University; Department of Biomedical Engineering, Northwestern University; Department of Physical Therapy and Human Movement Sciences, Northwestern University; Department of Radiology, Northwestern University; Department of Neuroscience, University of Minnesota; Department of Neuroscience, Northwestern University; Department of Physical Medicine & Rehabilitation, Northwestern University

## Abstract

Direct electrocortical stimulation (ECS) is a well-established brain mapping technique that helps achieve safe and effective resection of epileptic foci, tumors or vascular malformations. Recent studies using electrocorticography (ECoG) suggest that ECS’ exerted effects on brain sites are determined by the roles of those sites in larger networks. However, ECoG has limited spatial coverage. Here, we used functional magnetic resonance imaging from eighteen participants during performance of five language tasks to assess the functional network signatures of cortical sites defined as critical for speech and language by ECS. We found that critical sites causing speech arrests (SA) and language errors (LE) exhibited lower local and global connectivity than non-critical sites. LE sites showed greater connectivity across sub-networks (communities) than both non-critical and SA sites, indicating their role as connectors across functional networks. This connector profile of LE sites was most robust when considering network connectivity across the entire brain. Connector sites were concentrated primarily in temporal and inferior parietal cortices. Finally, these network features accurately predicted which sites were critical in machine learning models. These findings provide a preoperative framework for predicting cortical sites critical for speech and language function, which may ultimately help accelerate or improve brain mapping.

## INTRODUCTION

Speech and language are complex processes that require the recruitment of multiple brain regions and networks^1^. However, little is known regarding their precise network organization during active speech and language behaviors. In people with brain tumors or epilepsy undergoing resections of abnormal brain tissue, direct electrocortical stimulation (ECS) is commonly used to help neurosurgeons avoid resecting “eloquent” regions in the brain, i.e., sites that are “critical” for speech and language function, with the goal of reducing post-operative functional deficits^2–6^.

Despite its many decades of use, ECS has several limitations. Functional mapping via ECS has false positive and false negative outcomes^7,8^, and it can cause side effects including seizures or after-discharges. ECS mapping also requires an extensive amount of time and effort, either in the operating room during awake craniotomies or in the epilepsy monitoring unit, where it can take many hours, and therefore be exhausting for the patient and time-consuming for staff. Further, the neural and network patterns that functionally distinguish a critical cortical site from its non-critical neighbors during active speech and language engagement are still poorly understood. Improving this understanding could enable us to predict the criticality of cortical sites preoperatively. Thus, it would be beneficial to use complementary neuroimaging and recording techniques to guide and speed up mapping by reducing the potential number of areas needed to test with ECS and uncovering areas that might not have been suspected with ECS.

Historically, lesion studies have localized speech and language function to distinct cortical regions including Broca’s^9,10^ and Wernicke’s areas^11^. However, language-critical sites are often distributed across the cortex rather than strictly localizing to specific regions^12^. Moreover, language and speech tasks are produced by activity in widely distributed brain networks^1,13–16^. What then is special about the sites identified as critical using ECS? Currently, there is consensus neither on the role of these sites nor how ECS interacts with them. Studies using functional magnetic resonance imaging (fMRI) and electrocorticography (ECoG) have, shown some spatial overlap between ECS critical regions and activation in those areas during tasks or rest state (e.g., high gamma band in ECoG and blood oxygen dependent level [BOLD] signal in fMRI)^17–20^, but have also shown activation in many non-ECS critical areas.

These conflicting results suggest a need to assess ECS’ effects on language networks using network neuroscience (graph theory) techniques, rather than individual sites of activation. Recent work applying functional connectivity metrics to fMRI and ECoG data has begun to capture network-level effects of ECS beyond what activation alone can reveal; however, these studies have largely relied on simple connectivity measures such as coherence^21^ and mainly used resting state recordings^17–19,22–26^. The configuration of these networks likely differs between rest and active task engagement, even though the intrinsic structure of connections may be preserved^27–29^. Functional connections may be stronger during active language tasks than during resting state due to greater signal modulation during the task^30,31^, which may lead to improved discriminability between critical and non-critical sites for speech and language.

Graph theory, a powerful mathematical tool of studying connectivity^32^, has been scarcely used to predict ECS-defined language criticality. In brain networks, graphs consist of nodes (cortical or subcortical sites) and edges (structural connections or functional correlations between node pairs)^33–39^. Graph metrics applied to these networks can quantify connectivity properties spanning multiple scales—from local clustering among neighboring nodes, to global integration across the entire network, to interactions between functional subnetworks (communities, groups of correlated nodes). While two studies examined the potential of graph metrics derived from resting state ECoG and fMRI to predict the location of ECS-critical sites^21,40^. To our knowledge, no one has used graph metrics derived from task-based fMRI to predict critical sites.

Our group recently examined network properties of critical sites where ECS caused speech arrest or language errors using ECoG high-gamma band (70-150 Hz) activity during spoken word-reading^41^. Both speech arrest and language error sites exhibited significantly lower local and global connectivity than non-critical sites. Language error sites in particular acted as connectors across groups of correlated nodes (i.e., functional communities^42^ or modules). Further, these network metrics predicted the criticality of the sites with relatively high accuracy. However, ECoG arrays typically record from relatively small (regional) portions of the cortex, as they are limited by clinical monitoring necessity. In contrast, fMRI enables analysis of connectivity across the entire brain, revealing how language-critical sites interact not only within regions but also with remote brain regions outside of the traditional language network, albeit at a coarser spatiotemporal scale than ECoG. We therefore hypothesized that task-fMRI could provide additional insights into the role of ECS-critical sites in the global brain network.

Here, we investigated the network signatures of sites defined by ECS as critical for speech and language computed across the entire brain using fMRI during five language tasks. We examined measures of global, local, and inter-community connectivity derived from task-fMRI and demonstrated that critical cortical sites exhibited significantly lower local and global connectivity, and higher inter-community connectivity than non-critical sites. This suggested that their role as connectors was similar in global fMRI-defined networks to that seen in more regional ECoG-defined networks. We then showed that these fMRI network signatures accurately predicted the criticality of a cortical site. There was some variability in network metrics across language tasks, suggesting that the relative recruitment of these sites was modulated by the specific cognitive demands of the task. Lastly, we demonstrated how these graph metrics were distributed across the entire brain. Several brain regions exhibited connector-like metric profiles (high inter-community and low global/local connectivity), in particular temporal and perirolandic cortices. This highlights the temporal lobe as a key region in facilitating connections across speech and language subnetworks. It also suggests that task-based fMRI connectivity patterns may be used to enhance and facilitate functional brain mapping.

## RESULTS

### Participant info and demographics

We analyzed fMRIs from eighteen participants, including five with epilepsy and thirteen who underwent awake craniotomy for tumor resection. fMRI was recorded as participants performed five distinct language tasks: antonym generation, reading comprehension, rhyming, silent word generation and picture naming (Fig. 1C; see Methods). All participants were left-hemisphere dominant. Patients with epilepsy required ECoG monitoring for treatment of their seizures and patients with brain tumors had ECoG arrays placed on the cortex during the craniotomy. Participants had a mean (± SD) of 96.4 ± 29.3 electrode sites, with 11.3 ± 6.8 sites that were labeled as critical using ECS and 79.1 ± 30.3 sites that were tested and labeled as non-critical (NC) (Table 1; see Methods). The critical sites were further subdivided into language error (LE) and speech arrest (SA) sites (Fig. 1A, B). Speech arrest was defined as cessation of, slowed, or delayed speech output, while language errors were defined as anomias, paraphasias, or impaired comprehension. LE sites were mainly in perisylvian regions and middle temporal gyrus. SA sites were predominantly in motor areas such as the precentral gyrus, though some were also found in superior temporal and inferior frontal regions (Fig. 2A). On average, participants had 6.0 ± 5.5 LE and 5.3 ± 7.4 SA sites. Twelve participants had both LE and SA sites, three had only SA and three had only LE sites (Fig. 2B). Fifteen (83.3%) participants had electrode coverage in frontal lobe, seven (38.9%) in parietal lobe, fourteen (77.8%) in temporal lobe and one (5.6%) in occipital lobe. There were a total of 1627 electrode sites, out of which 203 were critical (108 LE, 95 SA) and 1424 NC. Participant information and demographics are illustrated in Table 1.

**Figure 1.**
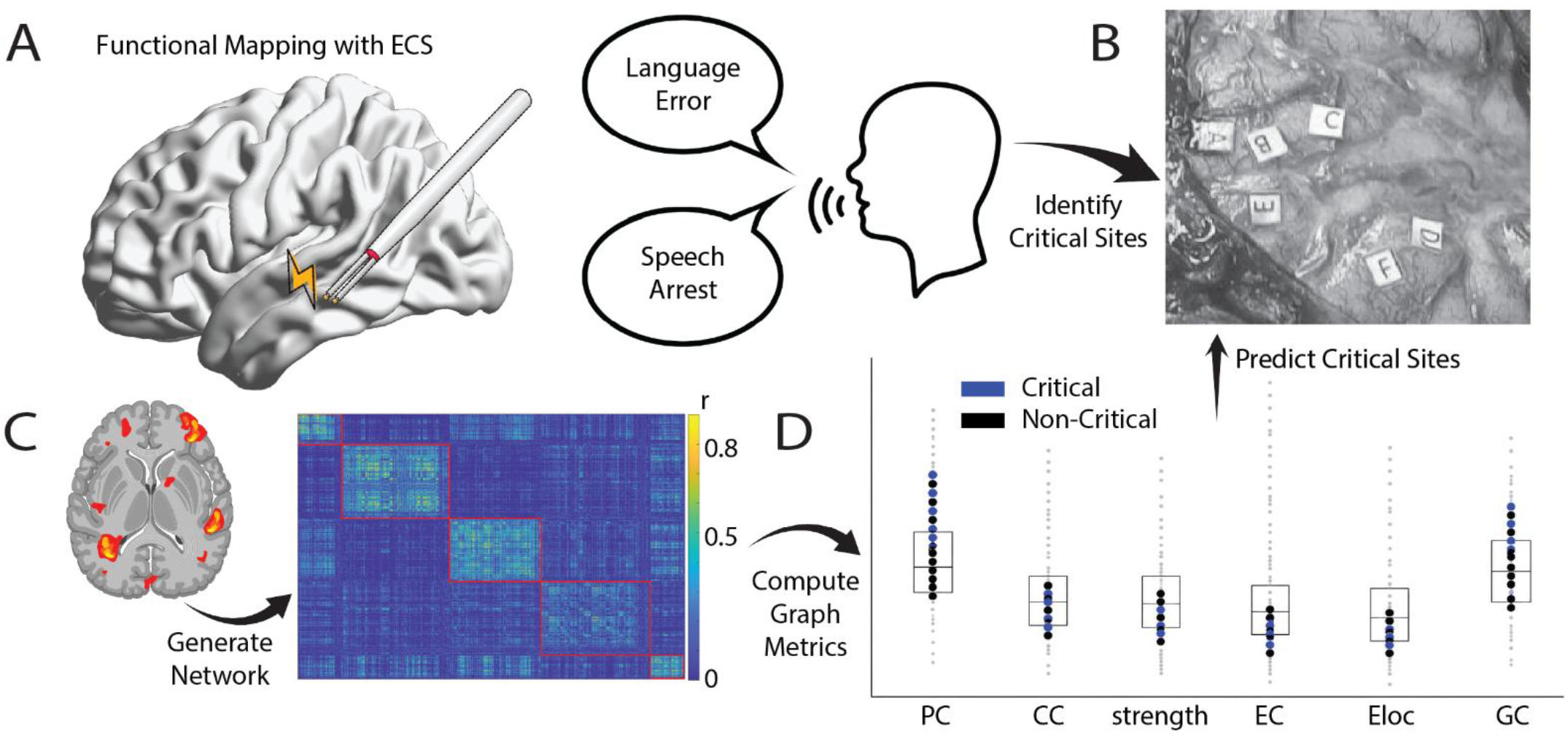
Methods summary. **A,** Functional brain mapping with ECS was performed intraoperatively (shown) for participants with brain tumors (n = 13) or extraoperatively in the epilepsy monitoring unit for participants with epilepsy (n = 5). ECS identified critical cortical sites by eliciting language errors (LE) or speech arrests (SA). **B,** LE and SA sites were labeled on the cortex following functional mapping. **C,** We recorded preoperative fMRI as the participants performed a subset of five language tasks: antonym recognition, picture naming, reading comprehension, rhyming and silent word generation. We generated a static network for each participant and task using pairwise BOLD signal correlations. The correlation matrix shown corresponds to one participant’s static network performing the antonym task filtered to include only positive correlations and reordered based on community assignments (shown with red outline). Communities were assigned using modularity maximization via the Louvain algorithm. The color bar indicates the Pearson correlation values (r) between each node in the network (Schaefer atlas and electrode ROIs). Brain activation figure on the left adapted from Biorender. **D,** We calculated six node-based graph metrics using the correlation matrices shown in **C**: participation coefficient (*PC*), clustering coefficient (*CC*), strength, eigenvector centrality (*EC*), local efficiency (*Eloc*) and gateway coefficient (*GC*). We then used these metrics as inputs to train machine learning classifiers (support vector machines and balanced random forests) using leave-one-participant-out cross-validation, to predict which sites would be critical to speech and language.

**Figure 2.**
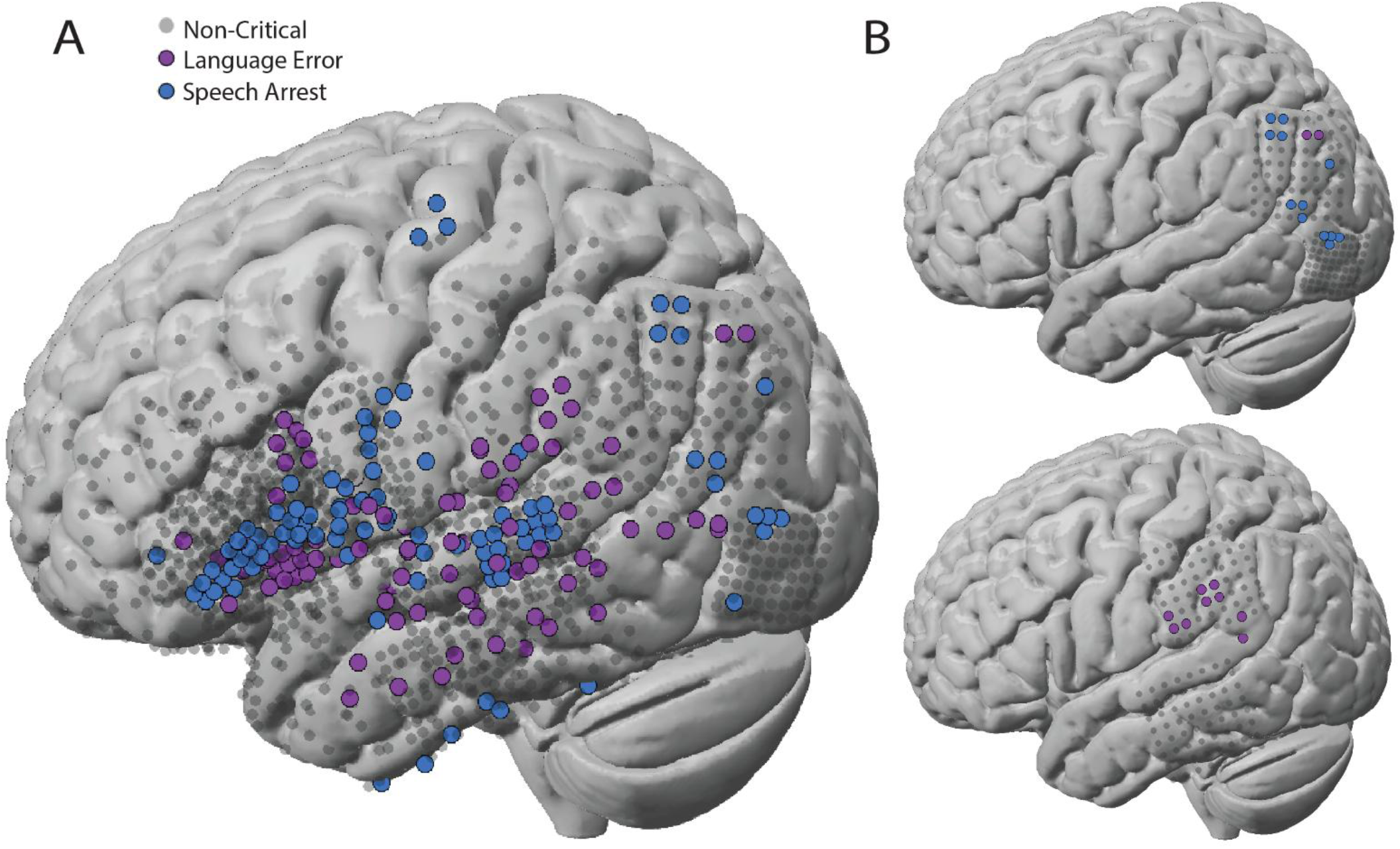
Electrode positions and example participant brains. **A,** All 1627 electrodes from 18 participants compiled and plotted on a template Montreal Neurological Institute (MNI) brain, with non-critical sites (n = 1424) in gray, language error (LE) critical sites (n = 108) in purple and speech arrest (SA) critical sites (n = 95) in blue. LE sites (in purple) were distributed across perisylvian regions— predominantly in ventral temporal and frontal regions but also located in posterior temporal and parietal regions. SA sites ( in blue) were mainly localized in premotor areas but were also widely distributed across ventral prefrontal and posterior and inferior temporal regions. **B,** Two example participant brain reconstructions (#11 and #10 from Table 1 on an MNI brain).

**Table 1.** Participant demographics and electrode information. ANT: antonym, PN: picture naming, RC: reading comprehension, RHYM: rhyming, SWG: silent word generation, R: right-handed, L: left-handed, A: ambidextrous.

| Participant | Type | Age | Sex | Tasks | Handedness | # of electrode sites | # of language error sites | # of speech arrest sites | # of non-critical sites |
| --- | --- | --- | --- | --- | --- | --- | --- | --- | --- |
| 1 | Epilepsy | 29 | F | ANT, RC | R | 69 | 8 | 6 | 55 |
| 2 | Tumor | 29 | F | All five | R | 46 | 7 | 2 | 37 |
| 3 | Epilepsy | 49 | F | PN, RC, SWG | R | 70 | 14 | 4 | 52 |
| 4 | Tumor | 40 | M | All five | R | 55 | 4 | 6 | 45 |
| 5 | Tumor | 61 | M | ANT, RC, RHYM, SWG | A | 61 | 4 | 4 | 53 |
| 6 | Epilepsy | 49 | M | ANT, RC | R | 63 | 2 | 8 | 53 |
| 7 | Tumor | 56 | M | All five | R | 80 | 3 | 1 | 76 |
| 8 | Tumor | 52 | F | All five | R | 66 | 4 | 2 | 60 |
| 9 | Tumor | 63 | M | All five | L | 129 | 15 | 1 | 113 |
| 10 | Tumor | 71 | F | All five | R | 129 | 9 | 0 | 120 |
| 11 | Tumor | 48 | M | All five | R | 128 | 2 | 12 | 114 |
| 12 | Tumor | 33 | M | All five | R | 130 | 0 | 32 | 98 |
| 13 | Tumor | 35 | M | All five | R | 67 | 0 | 11 | 56 |
| 14 | Tumor | 23 | M | All five | R | 132 | 0 | 4 | 128 |
| 15 | Tumor | 42 | F | ANT, RHYM, SWG | R | 129 | 2 | 1 | 126 |
| 16 | Tumor | 26 | F | ANT, SWG | R | 105 | 3 | 1 | 101 |
| 17 | Epilepsy | 11 | F | RC, RHYM, SWG | R | 70 | 13 | 0 | 57 |
| 18 | Epilepsy | 20 | M | RHYM, SWG | R | 98 | 18 | 0 | 80 |
| Total | 5 epilepsy, 13 tumor | - | 8F, 10M | ANT, PN, RC, RHYM, SWG | 16 R, 1 L, 1 A | 1627 | 108 | 95 | 1424 |
| Mean | - | 40.9 | - | - | - | 90.4 | 6.0 | 5.3 | 79.1 |

### Divergent functional network signatures delineate critical and non-critical sites

We defined the nodes of our network as regions of interest (ROIs) in two parts: 1) as spherical electrode sites with 2-mm radii, obtained from the spatial coordinates of each electrode in the ECoG array, and 2) as parcels from the Schaefer cortical atlas across the whole brain (1000-area parcellation, see Methods). We defined the edges of the network (a.k.a. functional connectivity) as the Pearson correlation coefficient of BOLD activity between all node pairs over the entire duration of the task (Fig. 1C). To investigate the functional connectivity patterns of critical and NC sites, we partitioned our networks into communities (subnetworks or modules) using modularity maximization via the Louvain algorithm and computed several well-known node-based network metrics (Fig. 3). We measured local connectivity with local efficiency (*Eloc*) and clustering coefficient (*CC*) (Fig. 3C), and global connectivity with node strength and eigenvector centrality (*EC*) (Fig. 3B). To measure each node’s inter-community connectivity, we calculated each node’s participation coefficient (*PC*) and gateway coefficient (*GC*) (Fig. 3A). We selected these graph metrics because 1) they allowed us to quantify a wide range of connectivity patterns from local to global to inter-network connectivity, and 2) our prior results using ECoG suggested that these metrics were indeed correlated with ECS criticality^41^. We hypothesized that critical sites would be more likely to bridge functional communities, i.e., act as connectors (Fig. 3A)—specifically, that high inter-community connectivity (*PC, GC*) and low local and global connectivity *(CC, Eloc, EC,* strength) would predict criticality. To test this hypothesis, we calculated each metric at each node (Schaefer ROIs and electrode sites) and compared the distributions of each metric for critical and non-critical sites using two-sample t-tests with false discovery rate (FDR) correction for multiple comparisons. To account for variability across tasks and participants, we standardized each metric by z-scoring within each participant and task prior to pooling all nodes for analysis.

**Figure 3.**
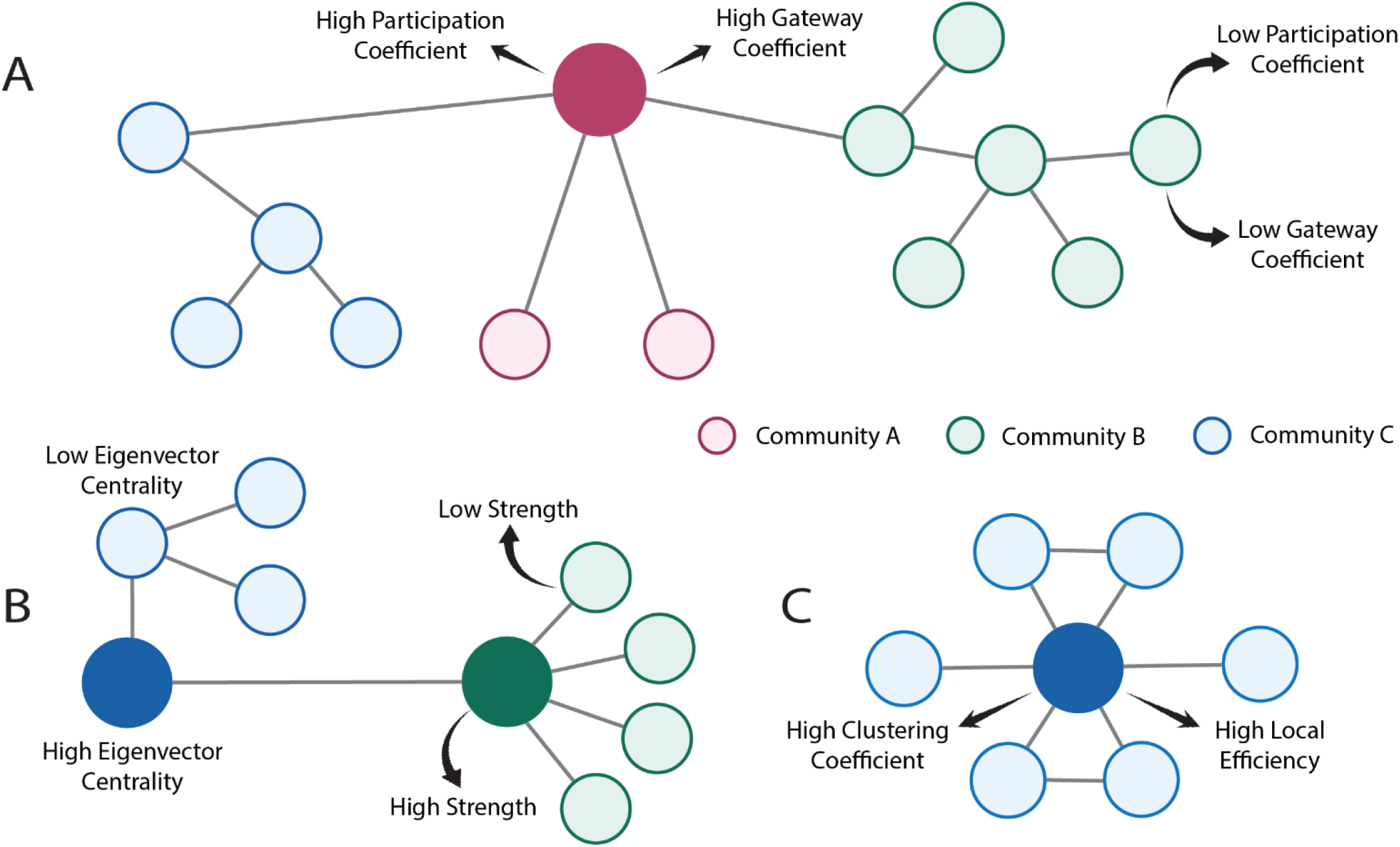
Diagrams illustrating the six network metrics. **A,** High participation coefficient and gateway coefficient characterize the large red node, as it is the only node in the network connecting across all communities (denoted in different colors). Nodes exhibiting this network profile are thus connectors. **B,** The large blue node has a high eigenvector centrality because its connections themselves have a high number of connections. Despite having the same number of connections as the large blue node, the smaller blue node has lower eigenvector centrality because its connections are connected to only one node in total (the large green node). A higher sum of connection weights in the large green node compared with the smaller green nodes results in a higher strength. High strength and eigenvector centrality nodes are typically global hubs, as they connect to many nodes within the network. **C,** The large blue node exhibits high clustering coefficient and local efficiency than the two small blue nodes on either side, due to its neighbors being more connected with each other (clustering) and a higher path length between its neighbors (local efficiency). This pattern is characteristic of local hubs, which cluster and connect densely within their own neighborhood.

We found that critical sites (LE and SA combined) exhibited distinct network signatures in the combined data across the tasks and participants. Critical sites exhibited significantly lower local (*CC*, *Eloc*) and global connectivity (*EC,* node strength) and higher inter-community connectivity (*PC, GC*) than NC sites (Fig. 4A and Table 2). Further, LE and SA sites exhibited distinct network signatures (Fig. 4B). SA sites exhibited significantly lower strength than NC sites, LE sites did not. LE and SA sites had significantly lower *Eloc* than NC sites, with LE showing significantly lower *Eloc* than SA sites. LE and SA sites exhibited significantly lower *CC* and *EC* than NC sites while there was no difference in *CC* and *EC* between LE and SA sites. Most notably, LE sites had significantly higher *PC* and *GC* than both SA and NC sites, suggesting that LE sites in particular act as connectors among functional subnetworks, rather than as local hubs, and that they tend to be the only connections across functional communities. This trend of high *PC* in LE sites was consistent across different community parameter thresholds (Supplemental Fig. 1). SA did not differ from NC nodes in *PC* and *GC*.

**Figure 4.**
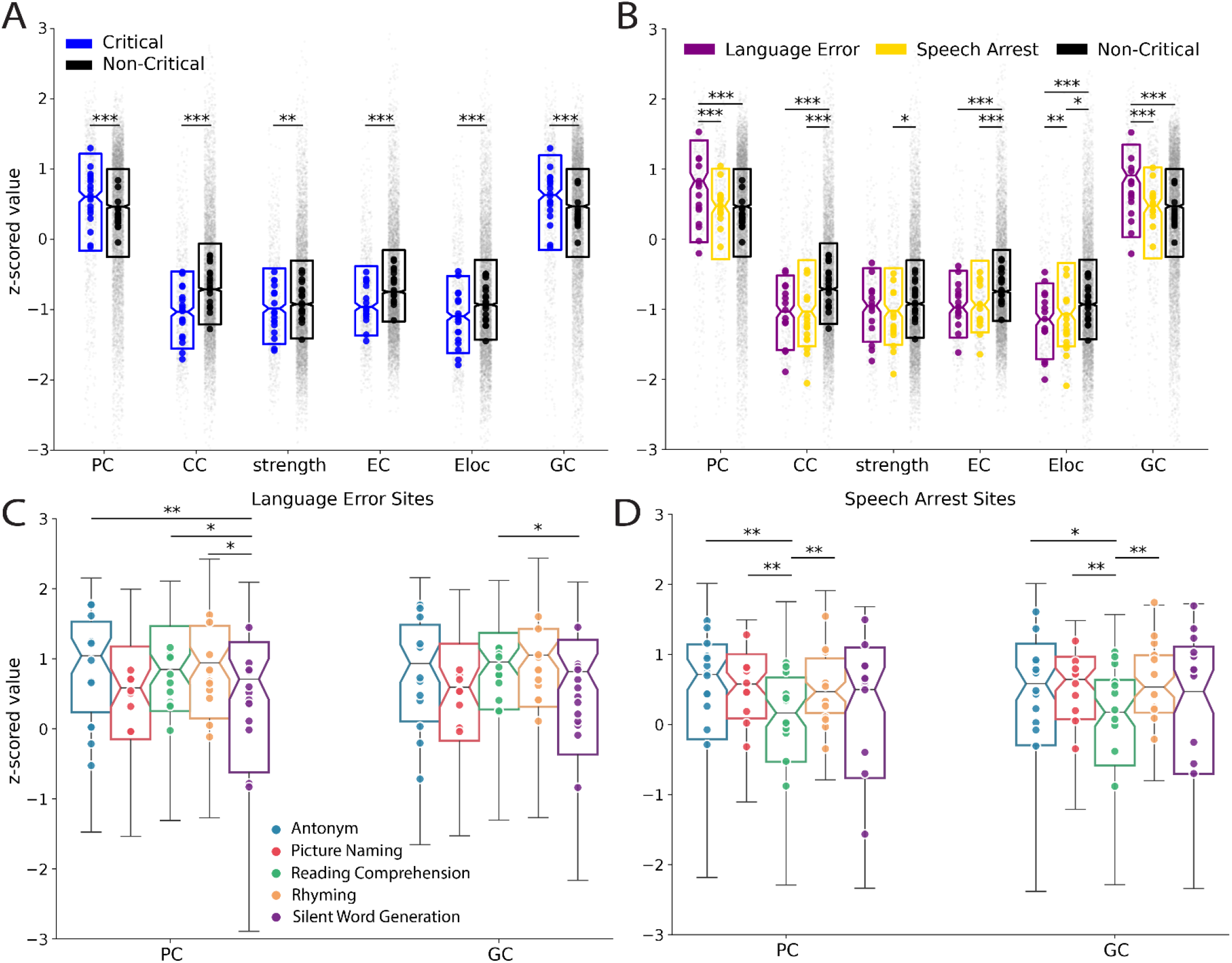
Network metrics. Distributions of participation coefficient (*PC)*, clustering coefficient (*CC)*, eigenvector centrality (*EC),* local efficiency (*Eloc)* and gateway coefficient (*GC)* across electrode sites in all five language tasks. **A,** Critical sites (n =203) exhibit significantly elevated *PC* and *GC* but lower *CC*, strength, *EC* and *Eloc* than non-critical (NC) sites (n = 1424). Critical sites include both language error (LE) (n = 108) and speech arrest (SA) (n = 95) sites combined. **B,** Metric distribution for LE and SA sites. High *PC* and *GC* in the critical sites are driven predominantly by the LE sites. *PC* and *GC* are significantly higher in LE sites than either SA or NC sites. *CC*, *EC* and *Eloc* are lower in both LE and SA sites than NC sites, while strength is only lower in the SA sites than LE or NC sites. All metrics were z-scored for each participant and task prior to plotting. Mean values of each participant are in bold, colored dots. Boxes indicate the median and IQR of each metric. While Schaefer ROIs were included to compute the network metrics, they are excluded from this figure to demonstrate the trends between our electrode sites. **C, D,** Values of participation coefficient (PC) and gateway coefficient (GC) in the LE sites **(C)** and SA sites **(D)** in different language tasks. LE sites had significantly higher PC in the antonym, reading comprehension and rhyming tasks than in the silent word generation task. Similarly, LE sites had significantly higher PC in the reading comprehension task than in the silent word generation task. SA sites showed the highest PC in the antonym task and higher GC in antonym, picture naming and rhyming than in reading comprehension. Statistical testing was calculated with a two-sided two-sampled t-test on z-scored metrics across all nodes with FDR correction: *p < 0.05. **p < 0.01. ***p < 0.001 (FDR-corrected).

**Table 2.**
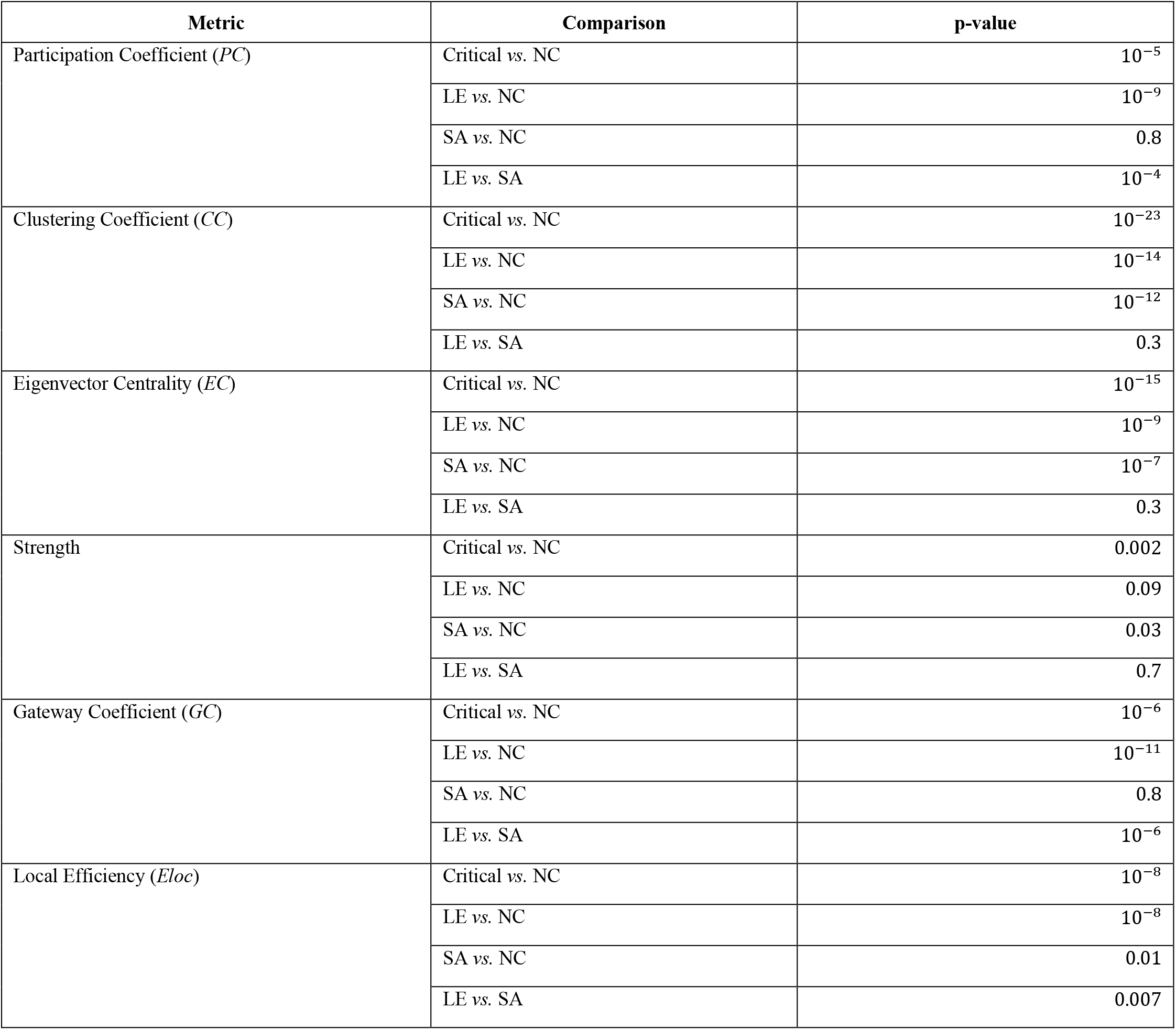
Summary of false discovery rate (FDR) corrected p-values for all metrics (two-sided t-tests, Benjamini-Hochberg).

| Metric | Comparison | p-value |
| --- | --- | --- |
| Participation Coefficient (*PC*) | Critical vs. NC | $10^{-5}$ |
| | LE vs. NC | $10^{-9}$ |
|  | SA vs. NC | 0.8 |
| | LE vs. SA | $10^{-4}$ |
| Clustering Coefficient (*CC*) | Critical vs. NC | $10^{-23}$ |
| | LE vs. NC | $10^{-14}$ |
| | SA vs. NC | $10^{-12}$ |
|  | LE vs. SA | 0.3 |
| Eigenvector Centrality (*EC*) | Critical vs. NC | $10^{-15}$ |
| | LE vs. NC | $10^{-9}$ |
| | SA vs. NC | $10^{-7}$ |
|  | LE vs. SA | 0.3 |
| Strength | Critical vs. NC | 0.002 |
|  | LE vs. NC | 0.09 |
|  | SA vs. NC | 0.03 |
|  | LE vs. SA | 0.7 |
| Gateway Coefficient (*GC*) | Critical vs. NC | $10^{-6}$ |
| | LE vs. NC | $10^{-11}$ |
|  | SA vs. NC | 0.8 |
| | LE vs. SA | $10^{-6}$ |
| Local Efficiency (*Eloc*) | Critical vs. NC | $10^{-8}$ |
| | LE vs. NC | $10^{-8}$ |
|  | SA vs. NC | 0.01 |
|  | LE vs. SA | 0.007 |

We further investigated how each graph metric varied across the five tasks. We hypothesized that distinct language tasks may differentially modulate functional connectivity patterns in critical *vs.* non-critical sites and thus lead to differences in graph metrics. All five tasks induced higher inter-community and lower local connectivity in critical than non-critical sites. However, there were some differences in these metrics across tasks. Notably, *PC* and *GC* in critical sites varied significantly across tasks (Fig. 4C, D). For example, *PC* was significantly greater in LE sites in the antonym, rhyming and reading comprehension tasks than in the silent word generation task (p = 0.01, 0.03 and 0.04 respectively; t-test, FDR-corrected). Similarly, *GC* was highest in LE sites during reading comprehension, during which it was significantly higher than during silent word generation (p = 0.04; t-test, FDR-corrected). In the SA sites, *PC* and *GC* were significantly lower during the reading comprehension task than during the antonym, picture naming and rhyming tasks (*PC*: p = 0.003, 0.01 and 0.006 respectively; *GC*: p = 0.02, 0.02 and 0.003 respectively; t-test, FDR-corrected). *CC*, *EC*, *Eloc* and strength did not differ across tasks for either type of critical site.

### Functional network signatures for LE sites are more apparent with global connections than regional connections

We next examined the effect of using smaller networks —i.e., within the regions covered by the ECoG arrays—in classifying critical sites. We computed our graph metrics on connectivity matrices computed using only parcels under electrodes, and not the Schaeffer ROIs from the rest of the brain. We found less difference in network metrics between critical and non-critical sites in smaller networks using only electrode sites (Fig. 5). Neither *PC* nor *GC* was significantly different between LE and NC sites (*p* = 0.3 and *p* = 0.6, respectively; t-test, FDR-corrected). The *CC* and *Eloc* remained significantly lower in LE sites than in NC sites (*p* = 10^−4^ and *p* = 0.005, respectively; t-test, FDR-corrected), but *EC* was no longer significant (*p* = 0.8; t-test, FDR-corrected). The *CC, EC,* node strength and *Eloc* remained significantly lower in SA sites than in NC sites (*CC*: *p* = 10^−10^, *EC*: *p* = 10^−9^, strength: *p* = 10^−9^, *Eloc*: *p* = 10^−10^; t-test, FDR-corrected).

**Figure 5.**
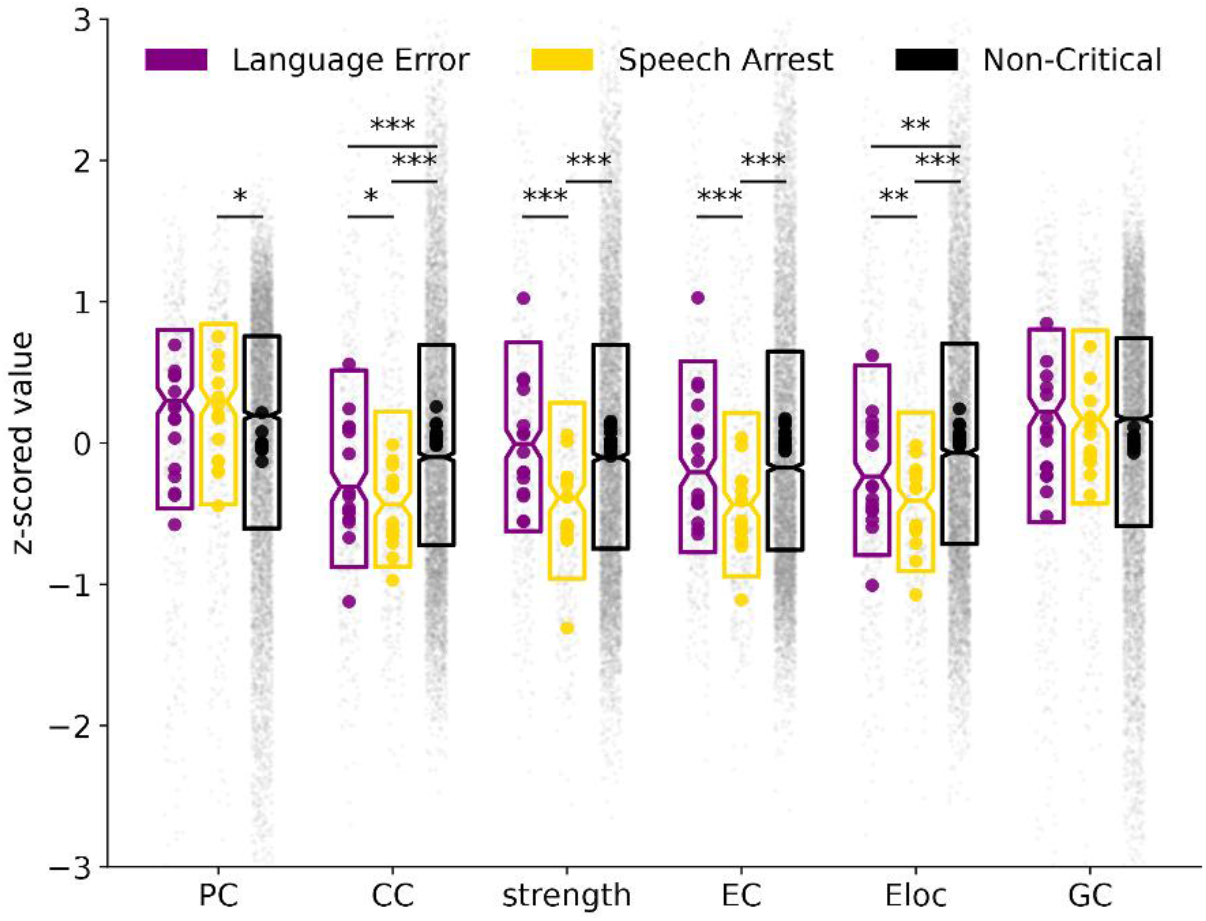
Network metrics computed using only local connections in all five language tasks. *PC*: participation coefficient, *CC*: clustering coefficient, *EC*: eigenvector centrality, *Eloc*: local efficiency, *GC*: gateway coefficient. No significant difference in *PC* and *GC* was found between language error (LE) sites and either speech arrest (SA) or non-critical (NC) sites when using only electrode ROIs. Statistical testing was calculated with a two-sided two-sampled t-test on z-scored metrics across all nodes with FDR correction: *p < 0.05. **p < 0.01. ***p < 0.001 (FDR-corrected).

### Measures of local and inter-community connectivity are distributed differently across the cortex

Critical sites tended to be located (Fig. 2) in perisylvian and temporal regions and displayed distinct network metric patterns, i.e., higher inter-community and lower local and global connectivity (Fig. 4A, B). Given these trends, we hypothesized that the anatomical distributions of these metrics across the whole brain could reveal the cortical regions that preferentially exhibit connector-like network organization during speech and language tasks (i.e., high inter-community connectivity and low local connectivity). We thus sought to examine the representation of these graph metrics across the whole brain, by projecting *PC* and *CC* within each Schaefer ROI onto a 3-dimensional brain within the Montreal Neurological Institute (MNI) coordinate system.

We found high values of *PC* in all lateral temporal regions, particularly anteriorly, and in the precentral, supramarginal, and angular gyri (Fig. 6A). *CC* was highest in the frontal lobe, particularly in parts of the left inferior frontal gyrus, and occipital lobe (Fig. 6B). Interestingly, *PC* and *CC* seemed to have inverse patterns of distribution across cortex. In the lateral temporal lobe, precentral and postcentral gyri, and supramarginal gyrus, *PC* values were high and *CC* values were low. While temporal cortices exhibited high *PC* in both hemispheres, the inferior frontal gyrus showed low *PC* and high *CC* only in the left hemisphere (Fig. 6A and 6B, top left). Further, *PC* and *CC* distributions showed substantial variability across tasks (Fig. 7). Picture naming and rhyming tended to show the broadest spread of high PC and low CC, especially in the dorsal stream (frontal and parietal areas). Lateral temporal areas consistently had high *PC* values in each task, especially anteriorly. *CC* was high in left inferior frontal gyrus and occipital lobe consistently across all tasks. The temporal lobe was consistently high in *PC* and low in *CC* across all tasks, which suggests that temporal areas are highly likely to be connectors across multiple functional communities during speech and language task engagement.

**Figure 6.**
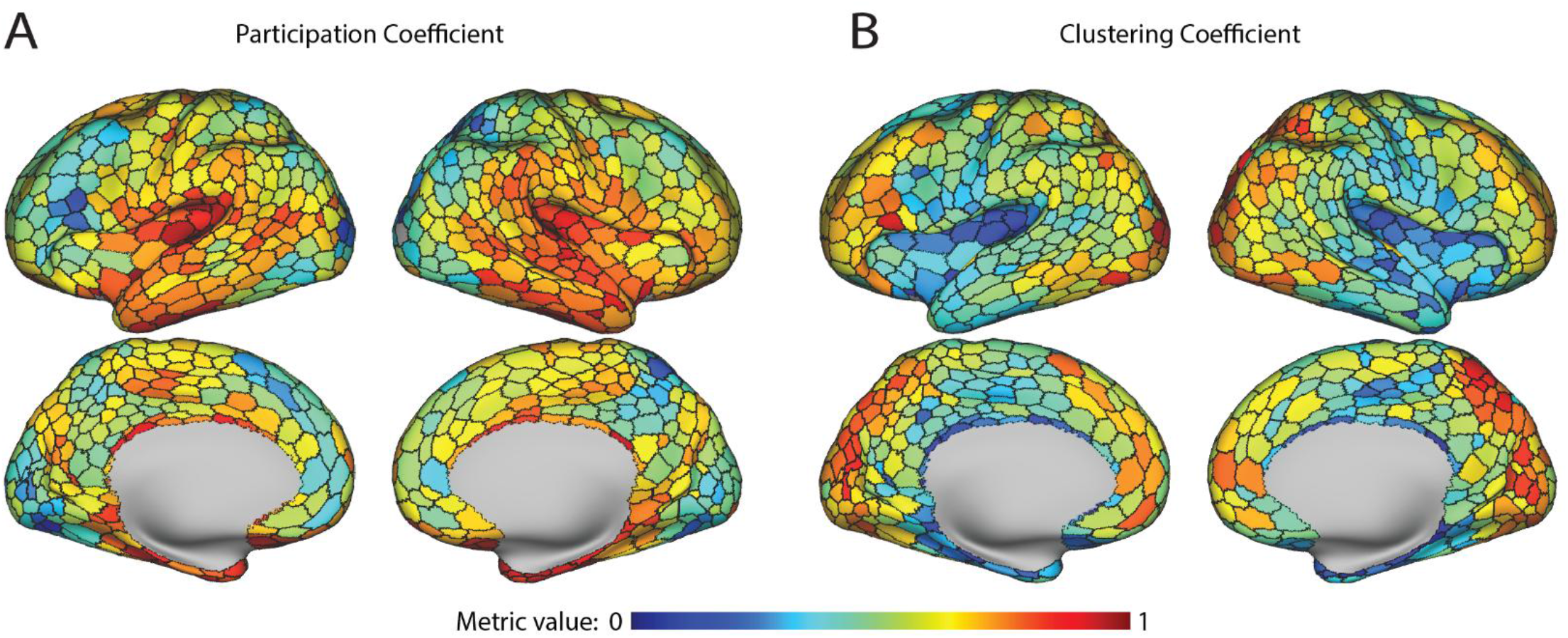
Anatomical distributions of network metrics. **A,** Participation coefficient (*PC*) and **B,** clustering coefficient (*CC*) across the whole brain. Both metrics in each parcel of the Schaefer cortical atlas were mapped onto an MNI brain via Connectome Workbench after scaling the metrics (min max normalization). The black boundaries correspond to the border of each Schaefer parcel. **A,** Premotor areas including the precentral gyrus, supramarginal gyrus, and temporal areas including most of the superior and some of the middle and inferior temporal gyri, and temporal pole exhibiting a notably high *PC*. In contrast, *PC* is lower in occipital regions and in the inferior frontal gyrus. **B,** *CC* exhibits a contrasting pattern to the *PC*, being especially high in the occipital lobe, inferior frontal gyrus and prefrontal areas. Pre and postcentral gyri and most of the temporal lobe exhibit low *CC*. While CC in temporal cortices were generally symmetrical in the left and right hemispheres, IFG was not; left IFG had high *CC*, while R IFG did not.

**Figure 7.**
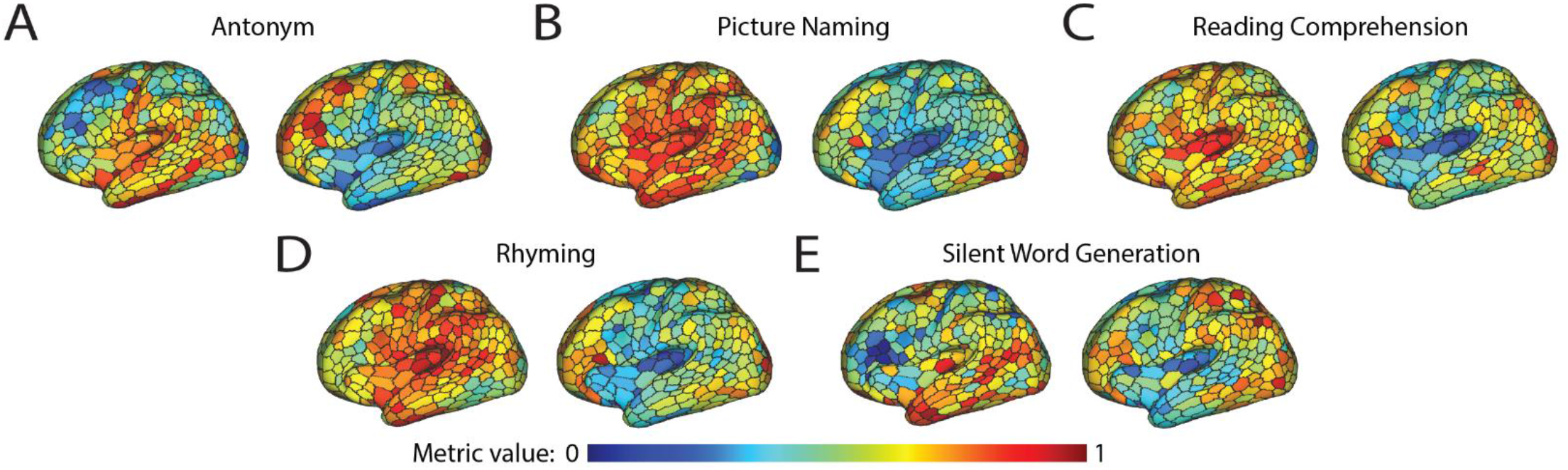
Task-level differences between network metric distributions. Participation coefficient (*PC*) and clustering coefficient (*CC*) in the antonym task **(A),** picture naming task **(B),** reading comprehension task **(C),** rhyming task **(D)** and silent word generation task **(E).** *PC* is shown on the left while *CC* is shown on the right for each subfigure. High *PC* and low *CC* values are distributed consistently across the temporal lobe, particularly anteriorly. The precentral, postcentral and supramarginal gyri exhibit high *PCs* in the antonym **(A),** picture naming **(B)** and rhyming tasks **(D).** Prefrontal regions exhibit higher *PC* distributions in the picture naming task **(B).** The left inferior frontal gyrus, mainly pars opercularis, has low *PC* in the silent word generation task **(E).** High *CC* values are predominantly in prefrontal regions, especially in the antonym, reading comprehension and rhyming tasks **(A, C, D).** Across all tasks, the occipital lobe and inferior frontal gyrus exhibit high *CC*.

### Critical sites can be predicted from their network metrics

We next assessed whether these network metrics derived from task-fMRI could be used to predict whether each cortical site was critical or not based on ECS. We trained support vector machines (SVM) and balanced random forests (BRF) to predict criticality in a binary fashion (all critical *vs.* non-critical, LE *vs.* non-critical, SA *vs.* non-critical). We trained the models using leave-one-participant-out cross-validation and compared our classifiers’ performance (balanced accuracy and sensitivity) to chance using permutation tests with 500 shuffles (see Methods).

Using combined data across all tasks, both SVM and BRF classifiers significantly outperformed chance level in both balanced accuracy (Fig. 8A) and sensitivity (Fig. 8B). Median balanced accuracy for classifying all critical sites combined *vs.* non-critical was 76.5% and 75.4% for SVM and BRF, respectively, which were both significantly higher than chance. Median sensitivity for classifying critical sites was also significantly above chance levels, at 77.1% and 64.0% respectively for SVM and BRF. Classification of LE and SA sites separately compared to non-critical sites was also significantly better than chance (Table 3). We predicted LE sites with 78.2% balanced accuracy and 70.0% sensitivity and SA sites with 87.8% balanced accuracy and 90.0% sensitivity. We achieved an AUC of 0.85 in our SVM predicting all critical sites, 0.92 in our SVM predicting SA sites and 0.83 in our SVM predicting LE sites (Fig. 8C).

**Figure 8.**
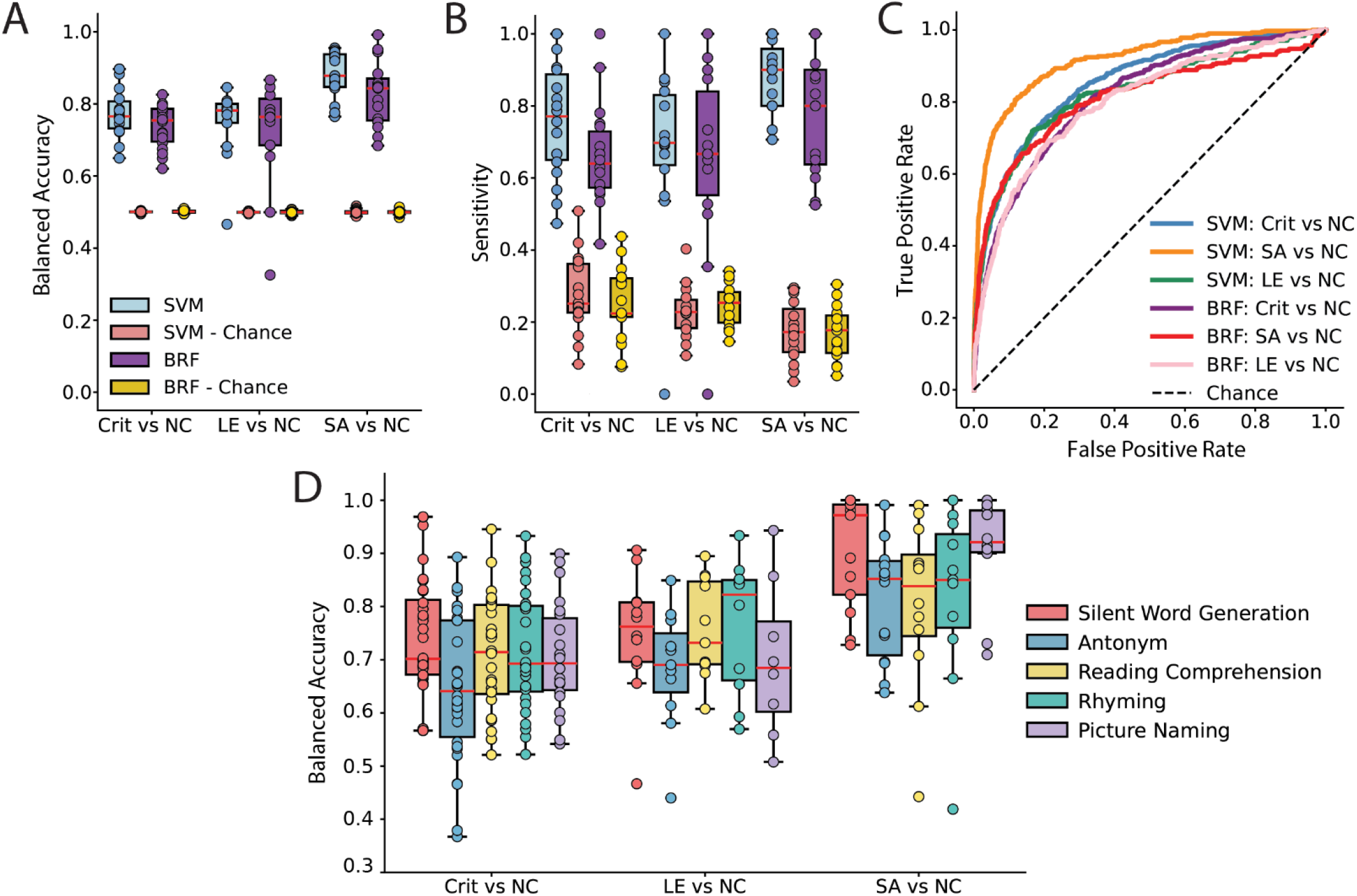
Balanced accuracy, sensitivity and ROC for critical site predictions. Support vector machine (SVM) and balanced random forest (BRF) classifier results in predicting critical sites in a binary fashion: Critical *vs*. Non-Critical (NC), Language Error (LE) *vs*. Non-Critical (NC) and Speech Arrest (SA) *vs*. Non-Critical (NC). **A, B,** Balanced accuracy and sensitivity in predicting critical sites significantly outperforms chance in all three binary models. All comparisons against chance are ***p < 0.001 (FDR-corrected). **C,** Receiver Operating Characteristic (ROC) curve represented for SVM and BRF models, with chance plotted as a dashed black line, with Area Under Curve (AUC) of 0.85, 0.92, 0.83, 0.81, 0.81 and 0.80 from top to bottom in legend. **D,** Balanced accuracy in each individual task as denoted by the legend, all significantly above chance levels at ***p < 0.001 (FDR-corrected). Differences in balanced accuracy between all task pairs did not reach significance.

**Table 3.** Summary of classification balanced accuracy and sensitivity. Median balanced accuracy and sensitivity values in support vector machine and balanced random forest models in predicting critical, language error and speech arrest sites. Median balanced accuracy and sensitivity averaged across all participants for both true labels (model) and permuted labels (chance) are compared with two-sided t-tests, corrected for multiple comparisons via Benjamini-Hochberg.

|  | Balanced Accuracy (%) |  |  |  | Sensitivity (%) |  |  |  |
| --- | --- | --- | --- | --- | --- | --- | --- | --- |
|  | SVM | Chance | BRF | Chance | SVM | Chance | BRF | Chance |
| Critical sites | 76.5 | 50.0 | 75.4 | 50.1 | 77.1 | 25.1 | 64.0 | 22.4 |
| Language error sites | 78.2 | 49.9 | 76.4 | 49.9 | 70.0 | 22.8 | 66.7 | 25.4 |
| Speech arrest sites | 87.8 | 49.9 | 84.3 | 49.9 | 90.0 | 17.3 | 80.0 | 17.7 |

**Table 4.** Summary of statistical testing for the decoding models. False discovery rate corrected p-values and area under the ROC (receiver operating characteristic) curve (AUC) between model *vs.* chance in predicting critical, language error and speech arrest sites. Median balanced accuracy and sensitivity averaged across all participants for both true labels (model) and permuted labels (chance) are compared with two-sided t-tests, corrected for multiple comparisons via Benjamini-Hochberg.

|  | p-values: SVM vs. Chance |  |  | p-values: BRF vs. Chance |  |  |
| --- | --- | --- | --- | --- | --- | --- |
|  | Balanced Accuracy | Sensitivity | Area Under ROC Curve | Balanced Accuracy | Sensitivity | Area Under ROC Curve |
| Critical sites | $10^{-18}$ | $10^{-12}$ | 0.85 | $10^{-18}$ | $10^{-12}$ | 0.81 |
| Language error sites | $10^{-10}$ | $10^{-7}$ | 0.83 | $10^{-6}$ | $10^{-6}$ | 0.80 |
| Speech arrest sites | $10^{-19}$ | $10^{-19}$ | 0.92 | $10^{-14}$ | $10^{-13}$ | 0.81 |

**Table 5.**
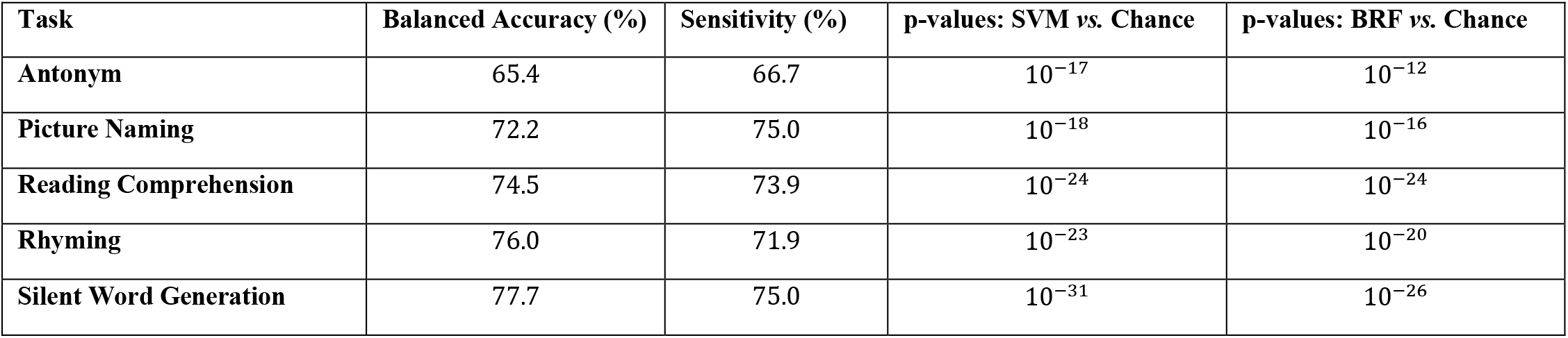
Summary of classification balanced accuracy, sensitivity and statistical testing for the decoding models across tasks. . False discovery rate corrected p-values between model vs. chance in predicting critical sites. Median balanced accuracy and sensitivity averaged across all participants for both true labels (model) and permuted labels (chance) are compared with two-sided t-tests, corrected for multiple comparisons via Benjamini-Hochberg.

We then trained our SVM and BRF separately on network metrics computed during individual tasks, again across participants. All classifiers performed better than chance in balanced accuracy and sensitivity, regardless of task (*p* < 10^−12^ in all cases, FDR-corrected t-tests). Balanced accuracy did not differ significantly among tasks for any of the binary classifications. However, there were some nonsignificant trends (Fig. 8D). For classifying all critical nodes, the antonym task tended to have lower accuracy than other tasks. In decoding SA vs NC, the silent word generation and picture naming tasks gave higher performance trends.

## DISCUSSION

It is not well understood what makes certain cortical sites in the brain’s entire network critical to speech and language function. Here, we identified functional brain network signatures underlying ECS-defined critical sites as well as how those signatures are anatomically distributed. Cortical sites identified as critical for either speech (SA) or language (LE) function exhibited low local and global connectivity, indicating that they are neither global nor local hubs. LE sites demonstrated high inter-community connectivity, indicating they act as connectors among functional brain subnetworks. These signatures of language-critical sites acting as inter-community “connectors” were most evident when our analysis included connections across the entire brain, rather than connections in only the regional language network. We observed this connector pattern most prominently in the anterior temporal lobe and inferior parietal lobule. Network metrics showed some variation among the different language tasks performed. Lastly, the network metrics enabled accurate classification of site criticality.

These results corroborate and extend our prior findings using ECoG^41^ that found the same network signatures in graph metrics between critical (LE and SA) and non-critical sites. Together, they provide strong evidence that connector nodes in the language network are more likely to be critical to language function. Language is a higher order process that requires interaction among multiple subnetworks, and our results imply that LE sites are important in coordinating activity among these subnetworks. Perturbing these sites with ECS may disrupt widespread information flow and thus lead to larger behavioral deficits. Further, given the lack of connector patterns but lower local and global connectivity in SA sites, their exact role in the speech and language network is uncertain and warrants further exploration.

Our ECoG recording study^41^ showed that LE sites were connectors within regional networks. In this study, LE sites were connectors, but only when including ROIs from the entire brain. This difference can be due to many factors. The tasks used in the two studies were different. Moreover, while BOLD activity has been shown to correlate with high-gamma activity^43^ (used in our prior study), BOLD is a much lower frequency signal, which may provide fundamentally different correlations within a network. Also, the smaller parcel sizes here likely had lower signal-to-noise ratios than high gamma power in our ECoG study, which could have affected correlations.

To our knowledge, no prior study has used graph metrics derived from task-fMRI to predict ECS-defined critical sites. Two studies investigated the correlation between resting state fMRI (rsfMRI) activity and critical sites, but did not use those correlations to predict critical sites^18,22^. Other reports found up to 1 cm (the width of an entire gyrus)^44^ and up to 2.5 cm^45^ between critical site locations and rsfMRI connectivity or task-fMRI-derived activation maps. However, distances between critical and non-critical sites often are just a few millimeters, warranting more exact localizations of criticality^12^. Our approach, by contrast, uses precise parcellations and graph metrics that shed light on the roles of critical sites in cortical networks.

We predicted which sites were critical on new, unseen data with reasonably high accuracy and sensitivity. While not currently accurate enough to be clinically viable, we anticipate this accuracy may improve with more participants’ data, more advanced network metrics, and the inclusion of other imaging modalities and/or ECoG. There are no comparable fMRI reports of accuracy in predicting ECS outcomes, but our results compare favorably with accuracies and sensitivities reported in prior studies using ECoG. In our recent study with ECoG^41^, we predicted critical sites with a median balanced accuracy of 65.9% across participants and 70.4% within participants. Two other studies using ECoG features to predict critical sites also reported high model performance^46,47^. Here, we predict critical sites with a median balanced accuracy of 76.5%, comparable to and even higher than performance metrics reported in prior studies. In addition, fMRI has one main advantage over ECoG: it can be performed preoperatively and thus is more easily scalable to clinical practice. This suggests these methods could pre-identify candidate critical sites before functional mapping and thus offer translational benefits in settings where ECoG is not available.

The anatomical distribution of graph metrics during speech and language function with a focus on ECS outcomes remains relatively unexplored. Many previous studies examining the spatial organization of functional hubs have primarily characterized regional differences using measures such as *PC*, degree, strength, and *CC*^15,48,49^. These studies have identified hubs in sensorimotor and parietal regions and high *PC* in regions including insula, inferior parietal sulcus, lateral occipito-temporal cortex, and superior parietal cortex. Another study with a patient cohort similar to ours used several fMRI-derived graph metrics to characterize regional network differences between low- and high-grade tumors, but focused on conventional, non-community-dependent metrics^50^. In this study, we investigated the spatial distribution of graph metrics that were most strongly associated with ECS-defined criticality: low local connectivity and high inter-community connectivity. In addition to elucidating how language tasks engage different brain regions, these anatomical distributions of graph metrics may also help guide functional mapping by pre-identifying regions with connector-like patterns in our participant cohort (i.e., regions more likely to be critical).

Across the entire brain, *PC* and *CC* exhibited nearly opposite patterns. Regions including the superior/middle/inferior temporal gyri, the temporal pole, precentral, postcentral, supramarginal and angular gyri exhibited high *PC* and low *CC,* indicating that they function as important connectors across subnetworks during speech and language processing. By contrast, parts of the prefrontal cortices and left inferior frontal gyrus exhibited low *PC* and high *CC*, suggesting specialized and modular roles. Our findings align with existing theories on the role of temporal pole, as well as inferior parietal lobule, as “hubs” for semantic or other language function^51–53^. Additionally, to our knowledge, previous studies have not explicitly reported a negative relationship between PC and CC across the whole brain. Thus, our findings suggest that these metrics capture distinct and inversely related aspects of network organization.

The left and right hemispheres also exhibited certain differences in metrics. The high local connectivity (*CC*) and low inter-community connectivity (*PC)* in the left inferior frontal gyrus, but not its right homolog, may reflect the left hemisphere’s specialized role in language processing. This hemispheric asymmetry may also be influenced by the task demands, given that *PC* was far lower in the left inferior frontal gyrus during the silent word generation task than the other tasks. Future studies incorporating a broader range of cognitive tasks alongside resting-state fMRI may help determine the extent to which these differences reflect task-dependent network organization versus intrinsic architecture.

The five behavioral tasks included here were designed to capture a range of linguistic processes for clinical purposes. Although decoding performance did not differ significantly across language tasks, several graph metrics exhibited task-dependent differences. For example, critical sites showed the highest *PC* during the antonym task. One plausible explanation is cognitive demand: the antonym task relies on multiple language subsystems and is more difficult than the other tasks. Task difficulty modulates brain activation patterns^54,55^, so this higher cognitive load have led to greater inter-community connections and thus higher *PC*. Identifying tasks that maximize these network-level distinctions might enable improved prediction of critical sites.

Our study had some limitations. Language tasks are inherently dynamic processes in which activity across multiple subnetworks rapidly fluctuates throughout task engagement. It is possible that the metric differences between critical and non-critical sites are heightened at distinct time points, which may be blurred with our static connectivity approach. Dynamic connectivity (of either ECoG or fMRI) elucidating how network features may fluctuate at distinct time windows may elucidate more nuanced patterns. Another limitation is the short duration of clinical fMRI scans, which may underlie the variability we observed across individual tasks. Further, our electrode parcels were designed to approximate the size of ECoG electrodes. Small parcels are inherently noisier than larger ROIs. While past studies have argued the interpretability of graph metrics decreases with the use of small parcels or voxels^49^, this specificity in defining the electrode sites was necessary to make direct comparisons across critical and non-critical sites within our ECoG arrays. Moreover, critical and noncritical sites often are within a few mm of each other, so the fact that we still obtained accurate prediction of critical sites and not at neighboring sites indicates that noise was not a major issue in these analyses.

Despite these limitations, our work demonstrates the functional network features that distinguish critical cortical sites from non-critical sites. Our ability to accurately classify site criticality in a diverse patient population highlights the potential for fMRI-derived network signatures to guide surgical mapping, reduce operative time, and ultimately improve the preservation of essential cognitive functions in patients requiring brain resections.

## METHODS

### Participants

In the present study, preoperative clinical fMRI was recorded on a total of 18 participants (age: mean = 40.9, range = 11-71, 8 females) who either required awake craniotomy and functional mapping for the resection of their brain tumors (n = 13) or had intractable epilepsy and thus required invasive ECoG monitoring (n = 5). All participants included in the study were native English speakers, left hemisphere dominant and had positive ECS cortical sites, i.e., sites where electrical stimulation led to speech arrest or language errors. Participants with any language or speech impairments, including but not limited to aphasia, dysarthria and apraxia of speech, were excluded from our study.

In participants with brain tumors, we placed 8×8 electrode arrays with either 3mm or 5mm interelectrode spacing over the frontal cortices and superior temporal and angular gyri, at least two gyri away from the tumor. In participants with epilepsy, standard clinical ECoG arrays and strips were placed with 1cm interelectrode spacing. All participants had ECoG coverage in the left hemisphere covering at least two of the frontal, parietal and temporal lobes. All participants gave written informed consent, in accordance with the Institutional Review Boards of either Northwestern University or Johns Hopkins University.

### Functional Mapping and Critical Node Definition

Participants underwent functional mapping using ECS as part of their awake craniotomy or for presurgical seizure localization. In participants with epilepsy, neighboring pairs of electrodes in the ECoG array were stimulated extraoperatively in the epilepsy monitoring unit. The two electrodes adjacent to a stimulation site were both deemed as critical if the site induced speech and language deficits upon stimulation. In the intraoperative setting, bipolar stimulation was delivered with an Ojemann stimulator after the patient was awake, starting at a current of 1mA and was increased in increments of 0.5mA until stimulation evoked deficits in speech and language. Tasks included picture naming, free speech, alphabet recounting and number counting. We designated all electrodes within a 5mm distance to a critical site as critical in the intraoperative setting. We then subdivided critical sites into two categories: speech arrest (SA) and language error (LE) sites. SA sites were defined as cortical areas where stimulation resulted in either consistent cessation of speech or slowed or delayed speech output while LE sites were designated as those causing any type of error in processing and understanding language, including but not limited to aphasia, dysnomia, and paraphasic and comprehension errors.

### MRI Acquisition

Functional and anatomical (T1-MPRAGE) MRI sequences were acquired on a 3T clinical scanner (Magnetom Verio, Siemens Medical Solutions USA, Inc. Malvern PA) at the MRI Facility of Northwestern Memorial Hospital, Chicago, IL, USA. fMRI imaging protocols consisted of a boxcar design with six cycles of alternating 20 seconds cycles of “task” and “rest” acquisitions. Each “task” block evaluated the response to 10 stimuli, with an interstimulus interval of 2,000 milliseconds (ms). Visual stimuli were presented on a mirror mounted on the head coil and that reflected an LED screen placed behind the scanner. Patients presented their response through a 2-buttons Lumina response pad. fMRI acquired 110 volumes during assessment of antonyms and 120 volumes for the other four language tasks. BOLD volumes were acquired with the following parameters: 220 mm FOV, 128×128 matrix, 31×3mm slices, and TR/TE=2000/20 ms. 3D MPRAGE was acquired for anatomical reference and BOLD overlay with these acquisition parameters 192×1mm thick axial slices, matrix = 256×256, FOV=256×256mm, TR/TE/TI/flip angle = 1800/2.55ms/9°. Following acquisition, MRI images were de-identified then stored on Northwestern University’s Quest High Performance Computing Cluster.

### Tasks

Participants performed several distinct fMRI tasks dependent on the location of the planned resection, with a total of five tasks: (1) an antonym task, (2) a silent word generation task, (3) a picture naming task, (4) a rhyming task and (5) a reading comprehension task. Out of all 18 participants, 15 performed the antonym task, 16 performed the silent word generation task, 11 performed the picture naming task, 14 performed the rhyming task, and 15 performed the reading comprehension task. 10 participants performed all five tasks.

#### Antonym

Participants were shown simple, non-emotional, commonly used English words and asked to think of a word with the opposite meaning. A plus (“+”) sign was shown during the rest phase. The scan consisted of five 22-second blocks per condition (task and rest), for a total scan duration of 220 seconds.

#### Silent Word Generation

Participants were shown upper-case letters (font size 200) and instructed to think of as many words as possible beginning with this letter. The scan consisted of six 20-second blocks, per condition (task and rest), for a total scan duration of 240 seconds. A meaningless non-letter symbol was presented as a baseline measurement during rest.

#### Picture Naming

Participants were shown pictures of everyday objects and asked to name the objects shown. Greek and mathematical symbols were shown during the rest phase of these two tasks. The scan consisted of six 20-second blocks per condition (task and rest), for a total scan duration of 240 seconds.

#### Rhyming

Pairs of words were shown on the screen and participants pressed one of two buttons on the Lumina pad when they believed that the words rhymed. During the rest phase, a pattern made from vertical or oblique slanting lines were shown and participants were required to press the button whenever the pattern matched. The scan consisted of six 20-second blocks per condition (task and rest), for a total scan duration of 240 seconds.

#### Reading Comprehension

A statement was displayed on the screen (font size 100), and participants were instructed to press any of the two buttons on the Lumina pad when they considered that the statement was correct. An array of Greek and mathematical symbols was illustrated during the rest phase. The scan consisted of six 20-second blocks per condition (task and rest), for a total scan duration of 240 seconds.

### fMRI Preprocessing

We analyzed all imaging data using Statistical Parametric Mapping (SPM12) in the MATLAB R2025b platform (The MathWorks Inc., Natick, MA, US), CONN (release 22.v2407), and custom MATLAB scripts. The first two initial volumes of each run were dummy scans to allow stabilization of the BOLD signal and steady state magnetization and thereby discarded. Following DICOM to NIFTI conversion, the remaining functional images were realigned to the first functional image to correct for head motion and resliced. Potential outlier scans with high motion artifacts were identified by calculating framewise displacement using the resulting six translational and rotational realignment parameters (framewise displacement values above 0.9mm were excluded^56^). The scans were then slice-time corrected (interleaved, bottom up). All functional and structural scans were co-registered using 4^th^ degree B-spine interpolation and segmented (FWHM: 60mm cutoff, number Gaussians: 1; repeated for 6 priors: grey matter, white matter, cerebrospinal fluid, bone, soft tissue, other). Functional and structural data were independently normalized into a standardized common coordinate system (Montreal Neurological Institute [MNI] space; IXI549Space). The resulting data were resampled to a common bounding box [-78 - 112 -70; 78 76 85] using 2mm isotropic voxels for the functional data and 1mm isotropic voxels for the structural data. Finally, all MNI-normalized functional images were spatially smoothed with a 4-mm FWHM Gaussian kernel.

### fMRI Processing

We extracted and denoised BOLD signal timeseries data in individual voxels using Ordinary Least Squares (OLS) linear regression, which projected each BOLD timeseries to the subspace orthogonal to all potential confounding variables, including noise artifacts from cerebral white matter and cerebrospinal areas, potential outlier scans and subject-motion parameters. Within the denoising step, subject-motion variability was minimized through the inclusion of 12 parameters, consisting of three translation and three rotation estimates alongside their respective first-order derivatives. We accounted for temporal autocorrelations and any physiological or head-motion sources using high-pass filtering with a set threshold of 0.01 Hz. Following denoising, we performed quality control of the timeseries data by visualizing the distribution of functional connectivity values between randomly selected pairs of voxels before and after denoising, to ensure properly centered distributions and reduced physiological and subject-motion effects. Finally, we performed ROI-to-ROI connectivity analysis using Pearson’s correlation coefficients on the extracted time series of each possible pair of ROIs (definition of ROIs and cortical atlas described below).

### Cortical Atlas and Definition of Individual ROIs

We defined the nodes of the network in two parts: (1) by parcellating each participant’s brain according to Schaefer’s cortical atlas^57^ consisting of 1,000 cortical nodes and (2) as spherical regions of interest (ROIs) computed from the MNI spatial coordinates of each electrode within each participant’s ECoG array, further labeled as critical and non-critical sites. We defined edges of the network as the BOLD activity cross-correlation between all node pairs over the entire duration of the task. As the electrode arrays only encompassed a small section of the brain, inclusion of a cortical atlas with a comprehensive coverage of the whole brain was necessary to investigate functional connectivity correlations between not just the electrodes but also between the electrodes and other brain areas outside of the ECoG array coverage. To localize the spatial coordinates of each participant’s electrodes in MNI space for intraoperative participants, we performed a neuroanatomical review of intraoperative photographs of the craniotomy before and after the placement of the ECoG array with the help of an expert neurosurgeon. Additional custom MATLAB scripts co-registered electrode arrays and ECS points to a three-dimensional MNI brain and extracted the xyz coordinate of each electrode site. For participants with epilepsy, we localized electrodes with a semi-automated pipeline by co-registering post-implant CT scans with preoperative thin-cut T1 MRI images, via LeGUI^58^. We then confirmed the locations of electrode arrays obtained from the automated pipeline by visualizing them on a three-dimensional MNI brain via custom MATLAB scripts.

We removed all voxels within the Schaefer atlas that overlapped with the electrode sites prior to fMRI processing via CONN to avoid duplicate voxels. ROI-to-ROI analysis in each participant and each task thus yielded a (1000 + # electrodes) x (1000 + # electrodes) functional connectivity matrix. Network metrics across all tasks were both combined and separated by task to assess any variations in connectivity and graph metrics in a language task-dependent manner. In the local connectivity section of our results where we computed graph metrics only on ROIs underlying electrode sites, our functional correlation matrices were in the form of (# electrodes) x (# electrodes), as we removed the 1000 Schaefer ROIs from this part of the analysis.

### Community Detection

We assigned communities using modularity maximization with the Louvain algorithm on our weighted correlation matrices, which creates the optimal community structure by maximizing the number of within-group edges and minimizing the number of between-group edges. The algorithm optimizes the modularity function 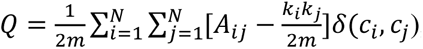, where *A_ij_* are the edge weights between nodes *i* and *j*, *k_i_*, *k_j_* are the sum of nodes *i* and *j*’s connections, *m* is the total sum of weights of all edges in the graph, *c_i_*, *c_j_* are the community assignments of nodes *i* and *j*, and *δ* is the Kronecker delta function. A greater value of *Q* corresponds to better partitioned networks of the graph, i.e., more within-community connections and fewer between community connections. The modularity quality function Q was optimized via consensus clustering, where the Louvain algorithm was repeated 100 times. The stability of these community assignments and metrics that depended on community assignments (participation and gateway coefficients) was tested by varying gamma, the community resolution parameter (Supplemental Fig. 1).

### Graph Metrics

#### Inter-Community Connectivity

We computed network signatures of the nodes in the network using the Brain Connectivity Toolbox (BCT)^59^. To quantify the inter-community connections of a given node, we calculated the participation coefficient (*PC*), defined as 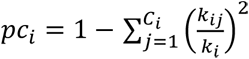, where *C_i_* is the community assignment of node *i*, *k_ij_* is the total number of connections of node *i* within community *j* and *k_i_* is the total number of connections of node *i* in the graph. Intuitively, the higher a node’s *PC*, the more connected it is to other functional communities within the graph. The *PC* is bounded by 0 and 1: if a node is only constricted to its own community, it will have a *PC* of 0 and if its connections are uniformly distributed across various communities, its *PC* will approach 1. Similarly, we assessed the inter-community connectivity patterns with gateway coefficient (*GC*), which is a variant of *PC* that is weighted by the number of inter-community connections of a node. In other words, a node will have a higher *GC* if it is the only connection between its community and another community. It is thus described as 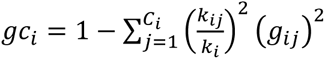, where 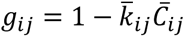.

#### Local and Global Connectivity

We quantified local connectivity patterns with clustering coefficient (*CC*) and local efficiency (*Eloc*). The clustering coefficient describes the proportion of the number of connections between a node *i* and its neighbors and the total number of possible connections between them. It is thus defined as 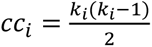. Local efficiency was calculated as the inverse of the distance matrix, which was calculated by converting edge weights to path lengths. We also investigated node strength, which is the sum of a given node’s total weighted connections, and eigenvector centrality (*EC*), defined as 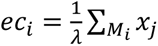 or simply *Ax* = *λx*, where *M_i_* is the set of neighbors of node *i* linked to node *j* and *λ* is the largest eigenvalue for matrix *A*. We only considered positive-strength edges in the graph when calculating these metrics, although their values when considering only negative-strength connections did not greatly differ than their positive-only versions. Each metric was z-scored for each participant and task and comparisons between LE, SA and NC nodes were conducted with false discovery rate corrected two-sample t-tests via Benjamini–Hochberg.

#### Whole Brain Projection of Graph Metrics via Connectome Workbench

Using the network metrics computed for each ROI within the Schaefer atlas, we projected *PC* and *CC* of all 1000 ROIs onto a 3-dimensional brain via Connectome Workbench^60^, resulting in a spatial distribution of *PC* and *CC* across the entire cortex. The electrode ROIs were excluded from the network in this analysis.

### Statistics

Prior to pooling metric data of all subjects and tasks, we z-scored network metrics individually within each participant and task and compared between critical and non-critical nodes using two-sided two-sample t-tests with false discovery rate (FDR) correction (Benjamini–Hochberg). Network metric values for all tasks were compared with two-sided two-sample t-tests between each task pair with FDR correction.

### Critical Node Prediction

We used support vector machines (SVM) and balanced random forests (BRF) to make three binary classifications for each node (ROI): critical *vs.* non-critical, language error *vs.* non-critical and speech arrest *vs.* non-critical. The SVM and BRF classifiers for our critical node predictions were trained using a leave-one-participant-out cross-validation paradigm, where the test set (one participant’s data) was left out of model training and hyperparameter tuning. For all classifiers, we excluded participants who did not have at least one node of the relevant class (i.e., at least one LE site for LE *vs.* NC classifiers, at least one SA site for SA *vs.* NC classifiers and at least one critical site (LE or SA) for Critical *vs.* NC classifiers). This resulted in 14 participants for the LE *vs.* NC classifier (4 participants excluded) and 15 participants for the SA *vs.* NC classifier (3 participants excluded) and no exclusions for the Critical *vs.* NC classifier. The training set (the remaining participants’ metrics) was used to train the classifier and optimize the hyperparameters of each model using 10-fold cross-validation for predicting critical sites and 5-fold cross-validation for predicting LE and SA sites. The 5-fold cross-validation for LE and SA sites was chosen to ensure that each cross-validation fold had at least one node of the relevant class. Parameters included the regularization parameter and kernel coefficient for the SVM, and the number of estimators and minimum split in each tree for the BRF. During cross-validation, we performed a grid search over a set of hyperparameter values and tested each separate training fold on the validation set, at the end of which the set of hyperparameters yielding the highest balanced accuracy on the validation set was chosen. Input features to all classifiers included the full set of graph metrics reported in our study: *PC, CC, EC, Eloc*, strength, *GC*. We also introduced penalty terms during model training that penalized misclassification of the minority class to reduce bias from class imbalance.

The average performance of the classifiers was compared to empiric chance by random label shuffling, which we repeated 500 times. We evaluated model performance using balanced accuracy, sensitivity, and receiver operating characteristic (ROC) analysis. For each classifier—SVM and BRF—we collected predicted decision scores and ground-truth labels across all cross-validation folds and concatenated them to form a single pooled evaluation set. This was done separately for each of three binary classification tasks: critical *vs* non-critical (Crit *vs* NC), speech arrest *vs* non-critical (SA *vs* NC), and language error *vs* non-critical (LE *vs* NC), yielding six ROC curves in total. To generate the ROC curves, we computed false positive rate and sensitivity across all classification thresholds and calculated the area under each ROC curve (AUC). We defined sensitivity as the true positive rate at each classifier’s default decision threshold (the sign of the decision function for the SVM; predicted class probability ≥ 0.5 for the BRF). All classification was done in Python using scikit-learn^61^.

---

**Supplemental Figure 1.**
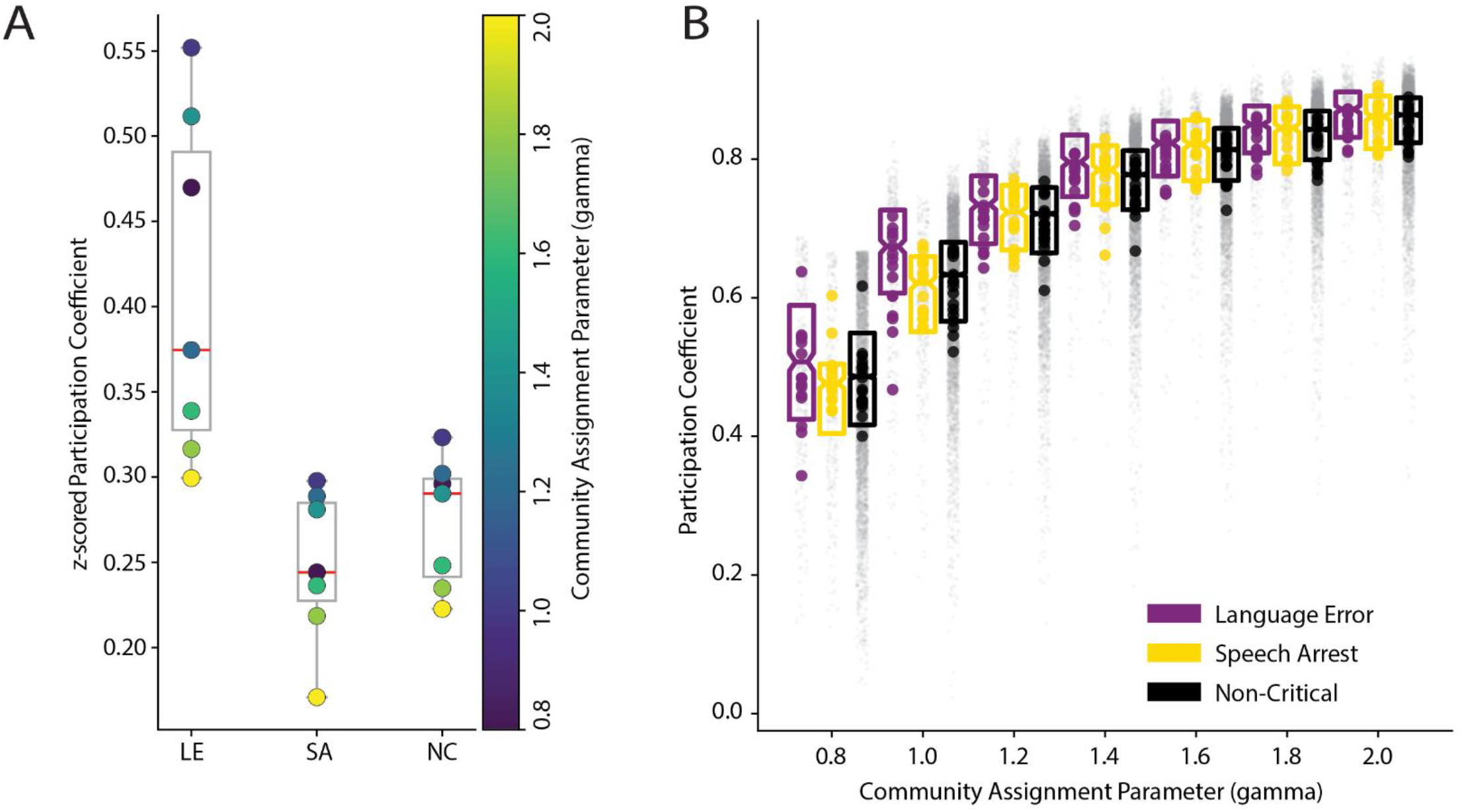
Stability of the participation coefficient (*PC*) relationships across site types. **A,** *PC* (z-scored) calculated at community assignment parameter (gamma) values ranging from 0.8 to 2.0. *PCs* computed below 0.8 gamma were excluded, as they resulted in only one to two communities and hence inconsistent PCs. Despite different gamma values, LE sites have consistently higher *PC* than either SA or NC sites. **B,** Raw *PC* values across different gamma values from 0.8 to 2.0. As gamma increases, so does the number of communities and subsequently *PC*. Gray datapoints are individual electrodes while bold colors represent participant averages.

## Contributions

**B.B**.: Conceptualization, methodology, software, formal analysis, investigation, and writing. **R.D.F**.: Methodology, software, investigation, and data curation. **N.E.C., M.C.T., M.A.M**.: Conceptualization, investigation, data curation. **Z.F., J.H.:** Methodology. **T.B.P, R.B.:** Conceptualization, methodology, and software. **M.W.S.:** Conceptualization, methodology, supervision, funding acquisition, and project administration. All authors reviewed and edited the manuscript.

